# *PaKman*: Scalable Assembly of Large Genomes on Distributed Memory Machines

**DOI:** 10.1101/523068

**Authors:** Priyanka Ghosh, Sriram Krishnamoorthy, Ananth Kalyanaraman

## Abstract

*De novo* genome assembly is a fundamental problem in the field of bioinformatics, that aims to assemble the DNA sequence of an unknown genome from numerous short DNA fragments (aka reads) obtained from it. With the advent of high-throughput sequencing technologies, billions of reads can be generated in a matter of hours, necessitating efficient parallelization of the assembly process. While multiple parallel solutions have been proposed in the past, conducting a large-scale assembly at scale remains a challenging problem because of the inherent complexities associated with data movement, and irregular access footprints of memory and I/O operations. In this paper, we present a novel algorithm, called *PaKman*, to address the problem of performing large-scale genome assemblies on a distributed memory parallel computer. Our approach focuses on improving performance through a combination of novel data structures and algorithmic strategies for reducing the communication and I/O footprint during the assembly process. *PaKman* presents a solution for the two most time-consuming phases in the full genome assembly pipeline, namely, *k-mer counting* and *contig generation*.A key aspect of our algorithm is its graph data structure, which comprises fat nodes (or what we call “macro-nodes”) that reduce the communication burden during contig generation. We present an extensive performance and qualitative evaluation of our algorithm, including comparisons to other state-of-the-art parallel assemblers. Our results demonstrate the ability to achieve near-linear speedups on up to 8K cores (tested); outperform state-of-the-art distributed memory and shared memory tools in performance while delivering comparable (if not better) quality; and reduce time to solution significantly. For instance, *PaKman* is able to generate a high-quality set of assembled contigs for complex genomes such as the human and wheat genomes in a matter of minutes on 8K cores.

## I. Introduction

*De novo* genome assembly is a fundamental problem in computational biology. The goal is to assemble the DNA sequence of an unknown (target) genome using the short fragments (called “reads”) obtained from it through sequencing technologies. The output is a set of “contigs” that represent contiguous portions of the target genome. Once assembled, the contigs are scaffolded, which is a step of ordering and orienting the contigs while accounting for potential gaps between successive contigs.

The genome assembly problem has been a topic of interest for well over three decades now, and yet the need for new scalable approaches has never been more critical than it is today. The factor driving this need is the continuously evolving landscape in DNA sequencing technology. With the advent of numerous high-throughput sequencing technologies, it has now become possible (even routine) to sequence a genome by running multiple clonal copies of the target genome (i.e., with *coverage C* ∈ [10, 100]), through a wetlab sequencing machine; and generating billions of reads (or hundreds of gigabytes to terabytes of raw data)—all in a matter of hours. For instance, a widely used technology such as Illumina is capable of generating short reads (~100 bases in length each) with an impressively low error rate (under 1%). There is also a new wave of “long read” technologies that are emerging in the community; however their error rates are still too high for wider adoption.

In this paper, we address the problem of generating a set of assembled contigs using short reads sequenced from an unknown target genome. (We do not consider the subsequent contig scaffolding step in this work.) There is a plethora of short read assemblers that have been developed for well over a decade (e.g., [1], [2], [3], [4], [5]). A large fraction of these assemblers use the *de Bruijn graph*, a graph data structure built out of fixed length substrings (of length *k*) contained in the reads, called *k*-mers, as their building blocks (vertices). Edges are established between vertices of any two consecutive *k*-mers in a read (overlapping in *k* − 1 positions). In contrast to older approaches, de Bruijn graph based approaches have demonstrated greater time-efficiency and high fidelity in genome reconstruction.

Despite their advantages, an efficient parallel implementation of a de Bruijn graph-based method on distributed memory platforms has proven to be challenging for various reasons. The input read set required to construct a de Bruijn graph is typically stored in the file system. In addition, many algorithms use the file system as additional space for intermediate memory-intensive algorithm phases. This, coupled with complicated I/O patterns, can potentially lead to an I/O bottleneck during the graph construction phase.

Secondly, to prune erroneous paths or prepare the output, a parallel implementation that uses a de Bruijn graph should be able to *manipulate* the graph in distributed memory post-construction. This is complicated by the inherently unstructured nature of the graph (as compared to, say, a dense matrix), leading to communication imbalance issues that necessitate specialized optimization strategies.

Finally, generating the output contigs involves performing numerous “walks” along the graph and enumerating the base pairs along the path as a contig. This creates multiple challenges under a distributed memory representation of the graph, requiring frequent coordination. For instance, one needs to ensure the same path is not repeatedly traversed by multiple processes to avoid over-representing a path in the contigs. This requires coordination (e.g., atomic operations) that could hamper parallel performance.

### Contributions

In this paper, we present the *PaKman* algorithm that addresses the above challenges. *PaKman* combines MPI I/O, MPI collectives, and a novel graph data structure (which we name *PaK-Graph*) to simplify I/O and communication patterns, and eliminate the need for expensive distributed-memory coordination during the walk phase. We evaluate *PaKman* on both shared and distributed memory platforms, demonstrating near-linear scaling behavior, and observe speedups between 1.6× and 3.2× over a state-of-the-art distributed memory assembler, and between 2.8× and 9.3× over state-of-the-art shared memory assemblers—all while producing comparable quality in output. To summarize, the key contributions are:

- A novel distributed memory data structure that enables contig enumeration with minimal coordination;
- A novel contig generation algorithm with simplified I/O and communication patterns; and
- Demonstration on shared and distributed memory systems.

## II. Overview of Approach

Fig. 1 summarizes the major steps of the proposed *PaKman* framework. Our approach to efficient and scalable genome assembly involves the following key optimizations:

**Fig. 1:**
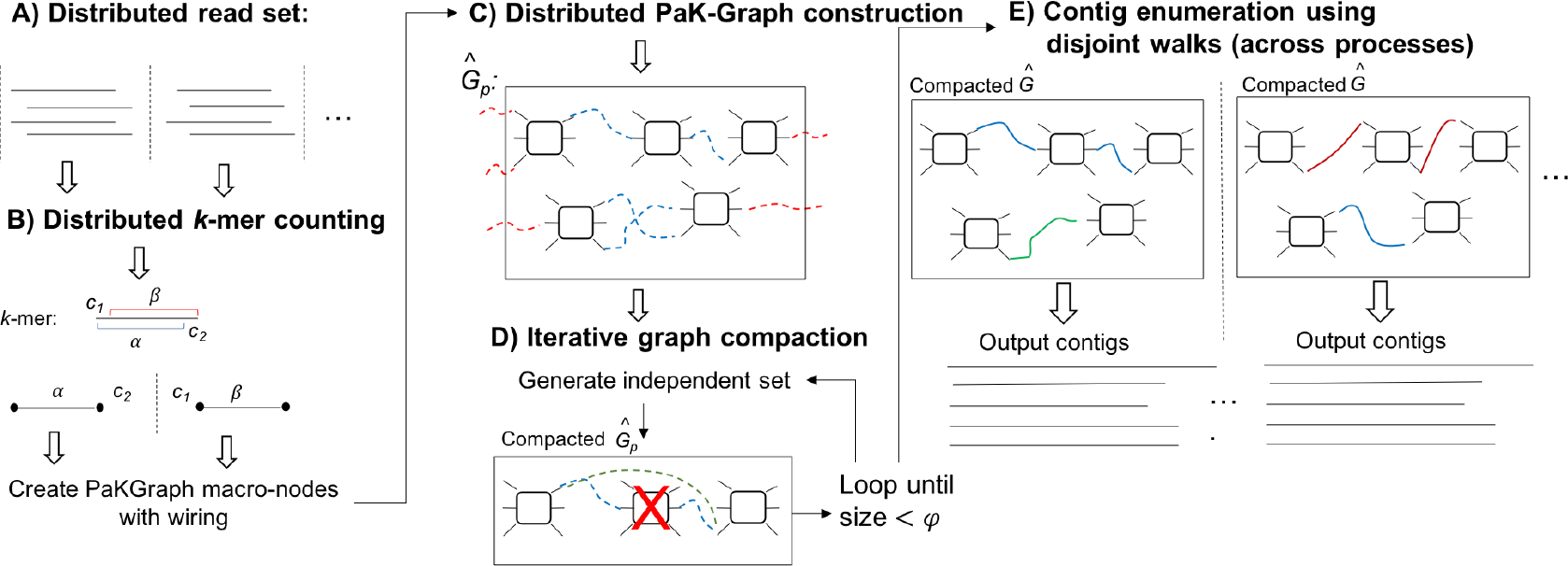
Schematic illustration of the *PaKman* assembly framework. Dotted arcs: edges in our PaK-Graph (shown only for illustration and not stored in our actual implementation); blue dotted arcs: edges internal to a process; and red dotted arcs cross process boundaries. Solid arcs: walks performed to generate contigs in parallel with replicated copies of the final compacted PaK-Graph as shown. 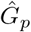: portion of PaK-Graph at process *p*, 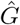: the global copy.

### Contiguous single-pass I/O

Each MPI process reads a distinct contiguous portion of the input file in parallel with other processes. After the input is processed in one pass through the file, no further I/O is performed. This minimal I/O requirement, coupled with the simple I/O pattern, makes it easy to efficiently utilize optimized I/O subsystems (parallel file systems, striping optimizations, burst buffers, etc.).

### Parallel load-balanced counting of k-mers

The input reads are processed to generate a stream of *k*-mersand the global count for each *k*-mer is computed. *PaKman* employs a scalable load-balanced algorithm to construct the *k*-mer histogram using only three types of MPI collective calls: `MPI_Allreduce`, `MPI_Alltoall`, and `MPI_Alltoallv`.

### Novel data representation and iterative compaction PaK-Graph

While input data and the de Bruijn graph might be space intensive, we observe that the final output of the algorithm is only of the order of Megabytes (or a few Gigabytes), depending on the species. This motivated the design of a compact representation of graph representing the *k*-mer connectivity. Rather than construct a conventional de Bruijn graph, we construct a new type of graph (which we call *PaK-Graph*; defined in Section III-B), which provides a way to arrive at a compact representation of the *k*-mers. This graph captures the *k*-mers and their connectivity in a lossless fashion. While the savings are not significant initially, we iteratively compact the graph further to dramatically reduce its size while preserving the total information. We demonstrate this compaction procedure can be performed efficiently in parallel.

### PaK-Graph replication and parallel deterministic walks

We compact the graph until it fits well within the memory available in each node of a distributed memory machine. We observe that the cost of compaction quickly decreases as the data structure shrinks in size, making subsequent compaction operations inexpensive. We exploit this property to sufficiently compact the graph to enable low-cost replication on all compute nodes of the parallel system. Once replicated, each MPI process picks a distinct set of starting points and performs disjoint walks to generate the contigs without any further communication or coordination. We achieve this using a deterministic algorithm to “wire” the paths through each vertex in a PaK-Graph to enable non-redundant walks without further coordination. This approach reduces the often complicated implementation of the walk phase into an embarrasingly parallel procedure (starting from each candidate *k*-mer) that incurs negligible time.

*PaKman* leverages algorithmic improvements to enable simplified communication and I/O strategies. Going further, the entire algorithm and its implementation rely only on a small number of MPI I/O and MPI collective operations, greatly simplifying performance portable implementations on new systems. The algorithm is efficient enough to outperform many shared-memory-specific de Bruijn based methods on shared memory platforms.

In the rest of the paper, we describe the design and implementation of each aforementioned step in detail.

## III. Methods

### A. Notation and Terminology

Let *r* denote a read of an arbitrary length (denoted by |*r*|) over the DNA alphabet *A* = {*a, c, g, t*}. For ease of exposition, we index the characters in a read from 1. Let *r*(*i*, *j*) denote the substring of length *j* starting at index *i* in *r*, such that *i* + *j* − 1 ≤ |*r*|. We denote the input set of *n* reads as 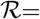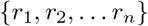, and their total length as *N* (= ∑_*i*_|*r*_*i*_|). We use · operator to denote string concatenation.

A *k-mer* in a read *r* is a substring of length *k* in *r*, for a given *k* > 0; similarly, a (*k*-1)-mer is a substring of length *k*-1. We use the term *l-mer* to denote a substring of length *l* that is significantly shorter than *k*—e.g., a typical value of *k* is between 32 and 48, whereas *l* is between 6 and 10.

### B. PaK-Graph: An Enhanced String Graph

In this section we introduce a new graph data structure called *PaK-Graph*, that we will use in our parallel algorithm (Section III-C). Given an input read set 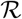, and positive integer constants *k* and *l* such that *l* < *k*, we define a directed graph 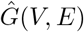 where *V* is the set of all “macro-nodes” and *E* is the set of all edges. We call each vertex in 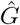 a “macro-node” because of its augmented node structure as defined below.

#### Macro-node

Each macro-node in 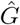 (as shown in Fig. 2a) is defined by a distinct (*k*-1)-mer present in 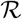. A macro-node *u* ∈ *V* has the following node structure:

- *label*(*u*) is the (*k*-1)-mer corresponding to *u*.
- *prefix_extensions*(*u*) is a set of arbitrarily long strings, each representing a candidate prefix extension of the (*k*-1)-mer (for an output contig).
- *suffix_extensions*(*u*) is a set of arbitrarily long strings, each representing a candidate suffix extension of (*k*-1)-mer (for an output contig).

**Fig. 2:**
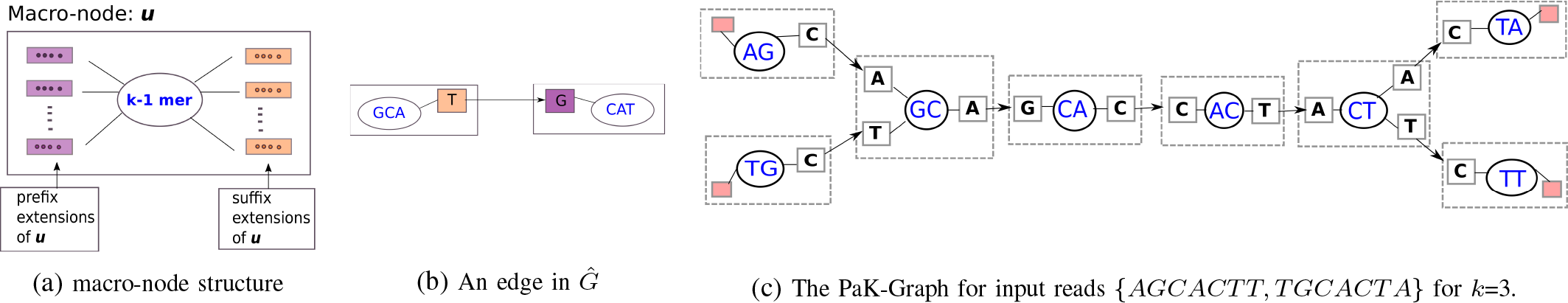
Macro-node structure and illustration

An extension with an empty string is called a *terminal* (prefix or suffix) extension.

#### Edges in 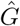

Edges in 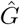 are defined between a suffix extension of one macro-node and a prefix extension of another. Specifically, there exists a directed edge *e* from a suffix extension *x* of a macro-node *u* to a prefix extension *y* of another macro-node *v* if and only if *label*(*u*)·*x* = *y·label*(*v*). Note that this implies there can be no more than one edge incident on each extension.

Fig. 2a represents a single macro-node identified by a (*k*-1)-mer. Fig. 2b presents two macro-nodes *GCA* and *CAT* connected by an edge such that, *GCA · T* = *G · CAT*. Fig. 2c presents an example of a PaK-Graph for a given input of two reads for *k*=3. The empty extenstions (shown in red) for the macro-nodes *AG*, *TG*, *TA*, and *TT* indicate there exists a terminal prefix extension for nodes *AG* and *TG* and a terminal suffix extension for the nodes *TA* and *TT*.

Initially, there exists one macro-node for every (*k*-1)-mer in 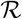. As for edges, for each *k*-mer *k*_1_ = *a*_1_*a*_2_…*a*_*k*_ in 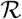, an edge is introduced in 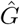 from the suffix extension with string *a*_*k*_ of the macro-node for *a*_1_*a*_2_…*a*_*k*−1_ to the prefix extension with string *a*_1_ of the macro-node for *a*_2_*a*_3_…*a*_*k*_. In this initial state, the PaK-Graph is equivalent to the de Bruijn graph constructed for 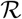. However, unlike the traditional de Bruijn graph, each macro-node through its extensions can encode an arbitrarily long path along the de Bruijn graph in a compressed manner. For this reason, our PaK-Graph can be viewed as an enhanced version of the string graph data structure originally introduced by Myers [6].

In the implementation, we only store the set of macro-nodes in 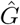. Edges are *not* explicitly stored because the suffix/prefix from the extension and the (*k*-1)-mer can be used to uniquely determine neighboring macro-nodes. Sections III-C3-III-C5 further detail the implementation of 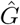.

### C. PaKman: Parallel Genome Assembly Algorithm

In this section, we describe *PaKman*, our parallel algorithm for genome assembly. The input is a set of *n* reads 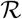 (made available as a single multi-sequence FASTA format file) and positive integer parameters *k* and *l* (*l < k*). The output is a set of *contigs* representing contiguous portions of the target genome. The number of processes is denoted by *psize*.

The algorithm consists of multiple steps as described below.

#### 1) Input Reading

The input is loaded from the input file in a distributed manner such that each process receives roughly the same amount of sequence data (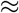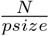 per process). This is achieved by each process performing a `MPI_File_get_size` and subsequently loading its unique chunk of reads using MPI-IO functions (`MPI_File_read_at_all`), such that no read is split among processes. Henceforth, we use 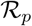 to denote the read set loaded at process *p* (i.e., 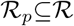).

#### 2) k-mer Counting

The goal of this stage is to generate and compute the frequency of all *k*-mers from 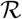. A well-known approach is to generate all *k*-mers by simply sliding a window of length *k* over each read and aggregating counts in a lookup table with 4^*k*^ buckets (one for each possible *k*-mer over the DNA alphabet) [7]. However, the large size of *k* (≥32) makes this simple approach prohibitive in space. Therefore, we use an alternative approach based on *minimizers* [8]. The idea is to use a smaller window length *l* (< *k*; e.g., *l* = 8) to partition *k*-mers into buckets, prior to obtaining the global count for each *k*-mer from each bucket. For parallel processing, each *min*-*lmer* bucket is assigned a distinct owner process. There are several ways to implement this minimizer approach using techniques from MinHashing based principles [9]. In our implementation, we assign a *k*-mer to the bucket corresponding to a least frequent *l*-mer occurring within that *k*-mer (i.e., making it the *k*-mer’s choice of its *min*-*lmer*). This way, we can expect (though not guarantee) that consecutive *k*-mers from the same overlapping region across reads are expected to be assigned to the same destination process bucket, which helps reduce communication later.

Algorithm 1 outlines our *k*-mer counting procedure. In the first step, each process generates all *l*-mers from its reads in 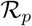 and obtains a global count for each *l*-mer using an `MPI_Allreduce` call. The next step implements the minimizer approach described above. This step involves redistributing the *k*-mers generated at various processes to their respective *min*-*lmer* buckets using `MPI_Alltoallv`. In our implementation, we perform this task using multiple rounds of communication to scale up to large input sizes (we use a batch size of a 100 million *k*-mers in all our experiments).

##### Algorithm 1 *k*-mer counting

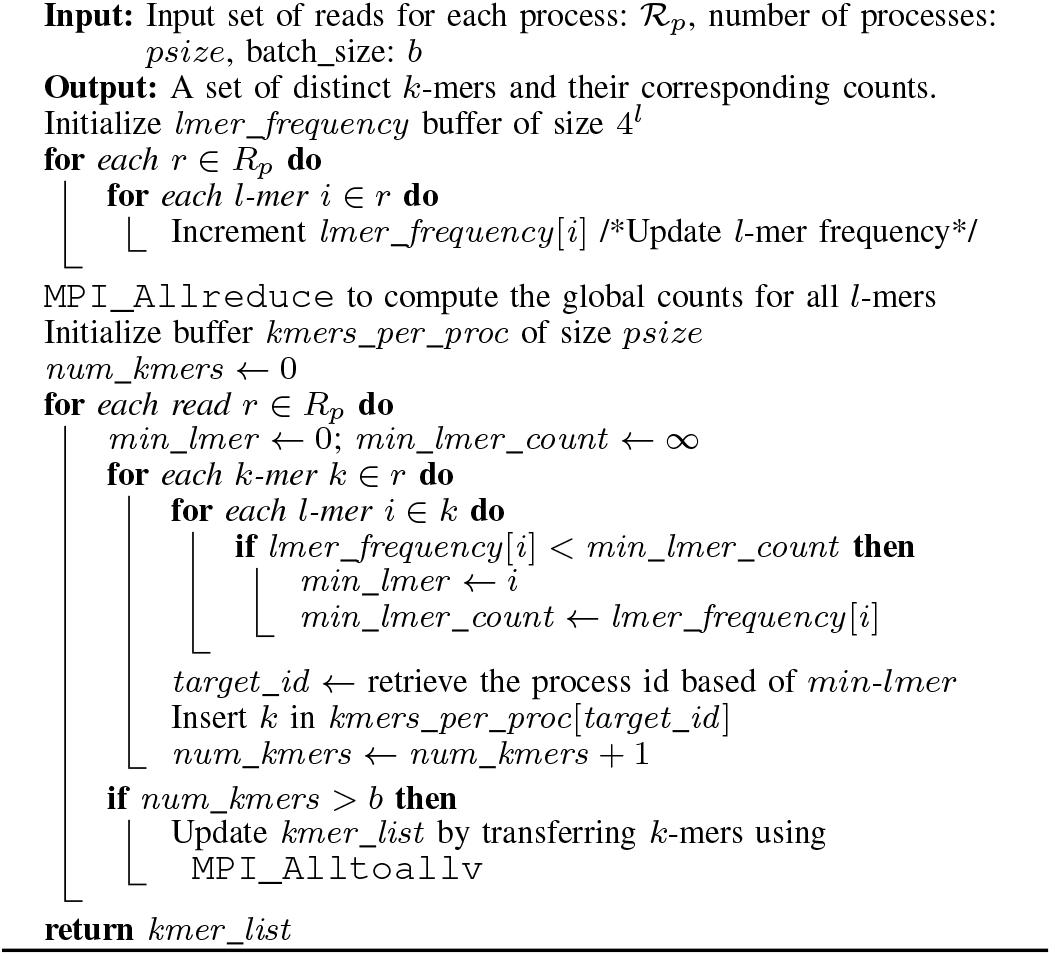

At the end of this step, all processes have a set of distinct *k*-mers and their respective global counts. Then, we perform a simple threshold-based pruning: we remove *k*-mers that have a count below a certain threshold *τ*. Such *k*-mers are deemed “poor quality” from the assembly perspective. We determine *τ* by plotting a *k*-mer frequency histogram for a fixed number of top buckets (say *h*)—obtaining the global counts using an `MPI_Allreduce`—and setting *τ* to the minimum over those *h* bucket counts. Parameter *h* is tunable and is set to 20 in our experiments.

#### 3) PaK-Graph Construction: k-mer Distribution

We now describe the distributed construction of the initial PaK-Graph, involving just a single `MPI_Alltoallv` communication. At the end of this step, each process *p* will hold a distinct portion 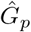 of the initial 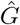.

Prior to constructing the PaK-Graph, we need to redistribute the *k*-mers because we need each *k*-mer in two places—one corresponding to the macro-node of its prefix (*k*−1)-mer and another to the suffix (*k*−1)-mer (as shown in Fig. 2). (If both (*k*−1)-mers are identical, then the *k*-mer is needed only in one place.) We identify the process id that will act as the destination for each macro-node using a linear congruential hash function for the macro-node’s corresponding (*k*−1)-mer; Subsequently, using an `MPI_Alltoallv` call, the set of *k*-mers are redistributed among the process space such that all *k*-mers corresponding a given macro-node are collected on a single process. At this point, each process *p* has a list of tuples 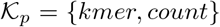 that will serve as the input to generate its 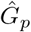.

##### Algorithm 2 Construct a PaK-Graph of macro-nodes

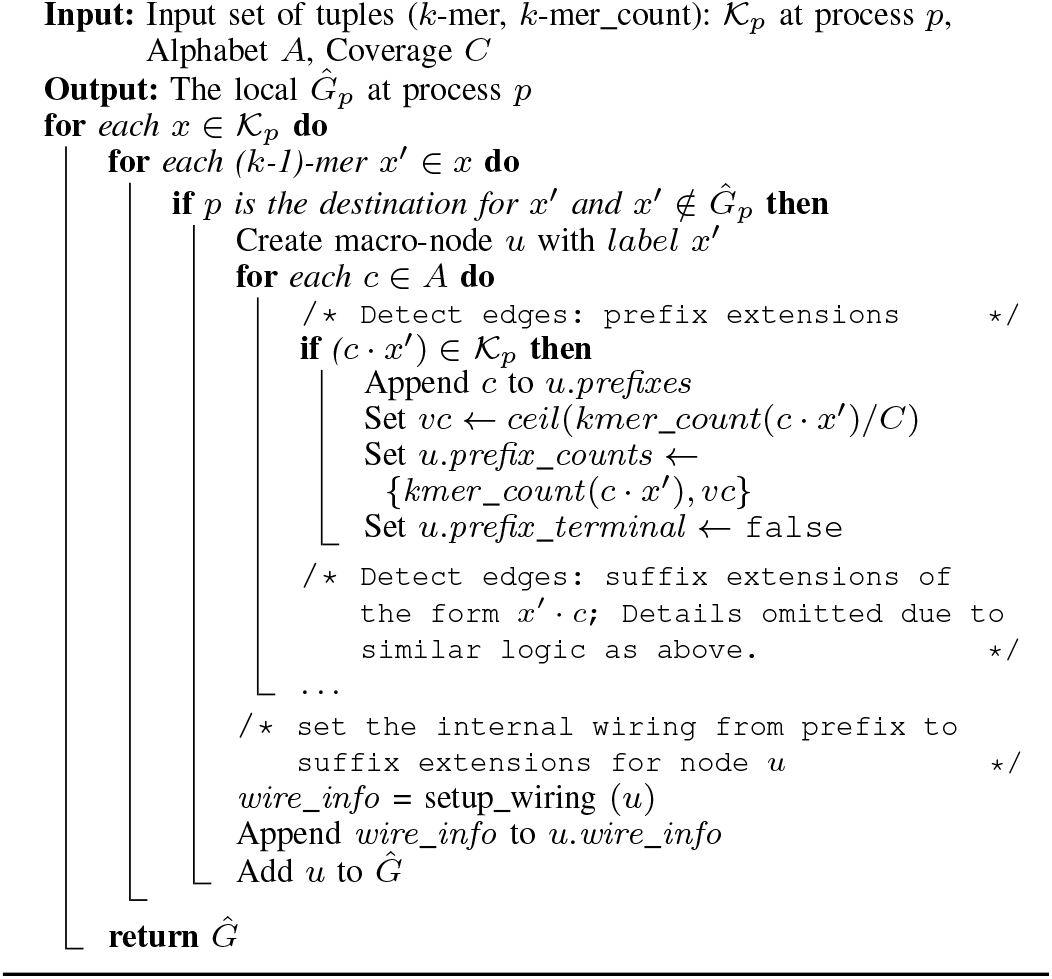

#### 4) PaK-Graph Construction: Macro-nodes

Algorithm 2 shows the steps to build 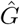 on each process *p* using 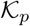. We make a couple of key observations here. First, a process *p* constructs a macro-node only if its *k*-mer falls in its domain (using the hash function). Second, as noted in Section III-B, the edges of 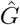 are *not* explicitly stored; instead, the extensions on either side of a macro-node are sufficient to capture all the information pertaining to its edges. However, how do we know if a particular extension exists or not (without communicating)? To answer this question, consider a valid prefix extension *c · x*′, where *c* ∈ *A* and *x*′ is the (*k*-1)-mer corresponding to the macro-node under construction. Then, *c* · *x*′ must be a *k*-mer that is also represented in 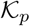 (as a result of the initial `MPI_Alltoallv`). This is the advantage of initially communicating *k*-mers to construct the local macro-nodes. In other words, the algorithm becomes communication-free at this step because all necessary information for macro-node construction is available from 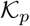.

There are two other steps in Algorithm 2 that need further elaboration. First, along with each extension, a list of pairs of the form 〈*kmer count, visit count*〉 is stored; where the *visit_count* represents the number of times that extension can be allowed to be traversed while taking part in contig enumeration (explained later). It is initialized to ⌈*kmer_count/C*⌉, where *C* is the sequencing coverage. At this time, we also determine whether a particular extension is terminal or not.

#### 5) PaK-Graph Construction: Wiring

Next, we compute a “wiring table” that holds the mapping from each prefix extension of the macro-node to a corresponding suffix extension. We explain the main idea of the wiring algorithm using the simple example in Fig. 3, which shows a macro-node for the (*k*-1)-mer *ACCT* (*k* = 5).

**Fig. 3:**
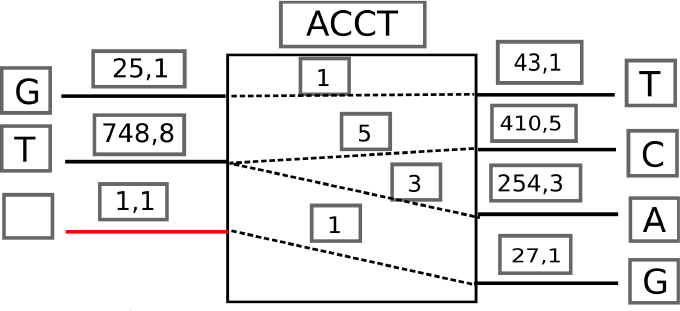
macro-node wiring illustration for (*k*−1)-mer *ACCT*. The pair <*kmer_count, visit_count>* labels each extension. The red prefix extension denotes a terminal prefix.

Initially, this macro-node contains two prefix and four suffix extensions (nonterminal), corresponding to a group of six *k*-mers. We first calculate the sum of all the prefix (*pc*) and suffix (*sc*) *visit_counts*. If the suffix (prefix) total count exceeds the prefix (suffix) total count, we introduce a new terminal extension on the prefix (suffix) side (shown in red), with the tuple 〈1, |*sc − pc*|〉 as shown. Subsequently, we construct a wiring table that connects each prefix extension to one or more suffix extensions (i.e., a fan-out). Note that multiple prefix extensions may also connect to one suffix extension (i.e., a fan-in). These wiring decisions are made based on the visit counts using a *greedy heuristic*. For instance, the prefix extension corresponding to *T* that has the maximum visit count (8) is considered first. This extension greedily selects the top available suffix extensions whose total visit counts become greater or equal to its own visit count—effectively selecting the suffixes *C* and *A* as shown. Ties are broken arbitrarily (albeit deterministically). This procedure is repeated until all extensions have been exhaustively wired.

Establishing a deterministic wiring strategy as described above helps us ensure that during traversal of the macro-nodes (in the contig generation phase), each walk is carried out in a coordination-free/disjoint manner—–instilling maximum concurrency in the process (Section III-C9).

This simple greedy strategy in wiring is also motivated by its impact on the quality of the output contig. Intuitively the (initial) visit count of an extension represents the number of distinct locations that particular *k*-mer (obtained by concatenating that extension with the (*k*-1)-mer) is expected to be present along the genome. Consequently, *k*-mers that are adjacent to this *k*-mer can also be expected (but not guaranteed) to occur with approximately the same frequency (e.g., if a *k*-mer ACCAG is present 10 times, then *k*-mers that represent one character extensions such as CCAGT or TACCA (if they exist) can also be expected to occur with similar frequency (without guarantee)). This is the intuitive reason behind the greedy strategy in wiring. We note here that our wiring strategy is amenable for extension to incorporate other qualitative information such as paired-end reads. If such information is made available, it could potentially have a larger positive impact on assembly quality. This is part of our future work.

#### 6) Contig Generation: Generate Independent Set

Using the initial 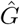, we initiate an iterative process of compacting the PaK-Graph until the total number of macro-nodes across all processes reduces to the extent that the entire graph will fit in the memory of each compute node. In our experiments, we set this threshold *ψ* to 100K macro-nodes.

Algorithm 3 describes the major steps of the iterative process. The main idea of the algorithm is to identify numerous macro-nodes for removal, remove them in a way that their information is captured in the macro-nodes that survive, and iterate with the compacted graph. In the interest of space, we omit the detailed algorithmic pseudocodes for the individual functions (*Generate_independent_set* and *serialize_and_transfer*); instead we summarize the main ideas in text alongside an illustration (Fig. 4).

**Fig. 4:**
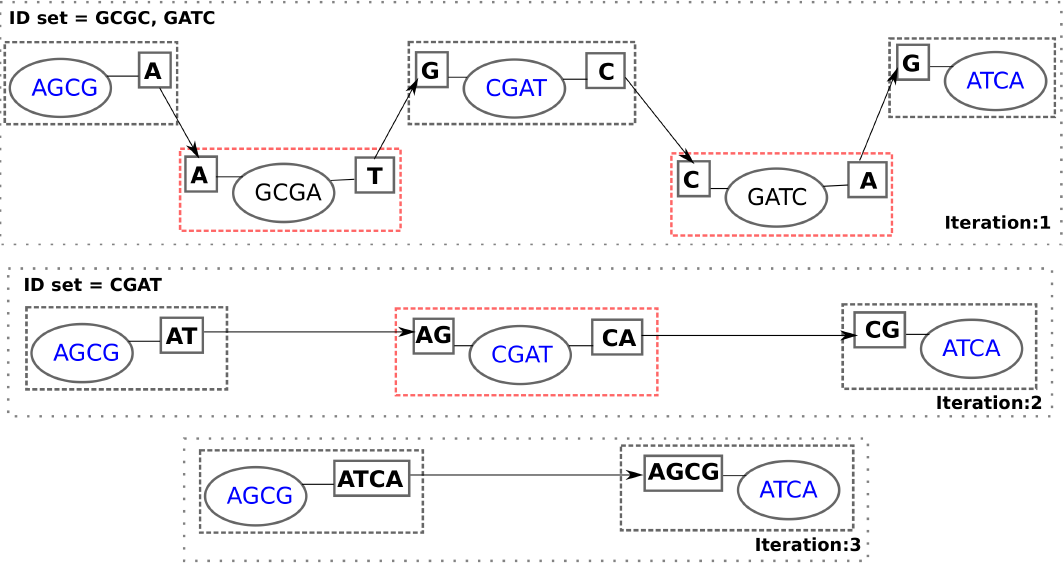
Illustration of iterative compaction of a five-macro-node PaK-Graph to a two-macro-node one in three iterations.

We formulate the problem of identifying macro-nodes to be removed as one of identifying an independent set *I* of macro-nodes in 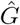. An independent set is a set of vertices in which no two are adjacent to one another. To identify an *I* corresponding to a 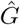, we use a simple distributed scheme in which each macro-node selects itself as part of the output set if and only if it contains the lexicographically largest (*k*-1)-mer among all its immediate neighbors. We devised this simple scheme because it enables each macro-node to make a strictly local decision without having to communicate with any of its neighbors. Surprisingly, we found this simple scheme also yields significant compaction. Specifically, in our experiments, we found the reduction in the number of macro-nodes between successive iterations ranged from ~25−28%, over the first few iterations. Such a sustained reduction would imply a 𝒪(log(*m/ψ*)) number of iterations required to converge, where *m* is the number of macro-nodes in the initial 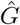.

##### Algorithm 3 Iterative algorithm to compact a PaK-Graph

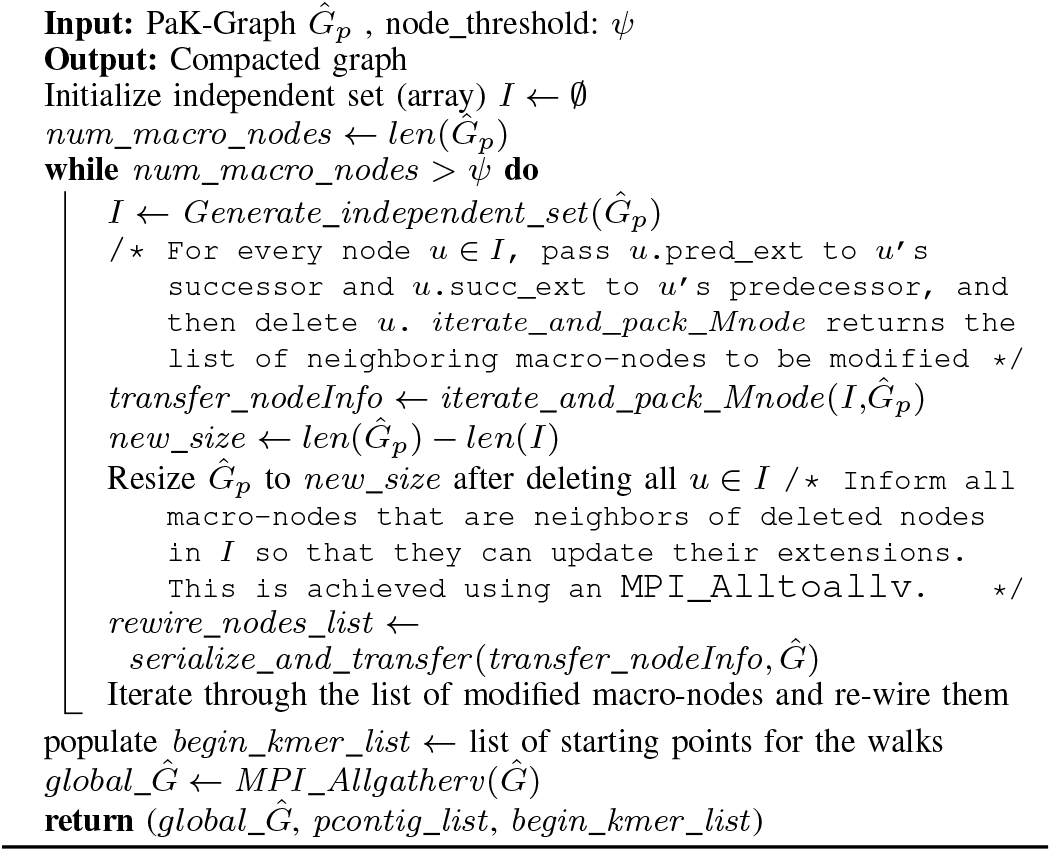

Intuitively, the motivation behind iterative compaction is to compress the graph to a state where the graph can be replicated in the local memory of each compute node and the contig enumeration step can be embarrassingly parallelized. The idea of detecting and using an independent set of macro-nodes to compact the graph at every step ensures that this compaction is achieved in a lossless manner. This is because no two macro-nodes to be removed at an iteration can be adjacent in an independent set, and a macro-node that survives this removal process carries forward the sequence information preserved in the corresponding extensions of the adjacent removed macro-nodes—as illustrated by the example in Fig. 4. This property holds true even in the more complex cases wherein a macro-node to be retained has multiple predecessor and successor macro-nodes (with one or more being part of the independent set); in such a case, our wiring scheme guarantees a deterministic pairing of all the added extensions at the surviving node, thereby resulting in no loss of data (proof omitted owing to space constraints). In terms of space complexity, with each compaction step, the removal of macro-nodes generates significant savings in practice, not only owing to the space constant (i.e., overhead) associated with each macro-node, but also by eliminating the redundancy that exists in the representation of *k*-mers among adjacent nodes in a PaK-Graph (i.e., the (*k*-1)-mer label and implicit edges).

#### 7) Contig Generation: Iterate and Pack nodes

In this step, the impact of removing the macro-nodes that are part of the independent set at each iteration is communicated to the surviving macro-nodes, so that they can update their structure (described in Section III-C8). In the first iteration of the example in Fig. 4, once the macro-node corresponding to label *GCGA* is removed, its two corresponding wired prefix-suffix extension pair, namely the macro-nodes for *AGCG* and *CGAT*, need to be informed. If any of these macro-nodes are remote, then the information about this deleted macro-node needs to be communicated. The *iterate and pack* function prepares the data to be communicated and the next step (*serialize and transfer*) performs the communication and macro-node update.

#### 8) Contig Generation: (de)Serialize and Transfer

In this step for iterative graph compaction, an `MPI_Alltoallv` communication call relays all removed macro-node information to the impacted processes. Subsequently, macro-node information at the impacted processes is updated based on the macro-nodes removed from 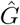. Consider again the example in Fig. 4. After removal of the macro-node *GCGA* in iteration 1, the neighboring macro-nodes now become immediate predecessor and successor (akin to the removal of a node in a linked list). Note that such new predecessor-successor relationship is established only between pairs of prefix-suffix extensions that have an entry in the wiring table of the macro-node being removed. As a result of this repacking, the suffix and prefix extensions of the two macro-nodes (*AGCG* and *CGAT*, respectively) should be extended as shown to include the values from removed macro-node. Note that if both a prefix and suffix extension wired pair for the removed macro-node happen to be terminals, then we construct and output the corresponding contig.

It is to be noted that the sizes of the macro-nodes in the buffer *transfer_nodeInfo* do not stay constant, owing to the varying lengths of the tuple entries that get communicated. In fact, the extensions tend to grow in size as the number of iterations grows. As a result, we need to serialize the contents of the *transfer_nodeInfo* buffer to convert it to a byte stream. We utilize *cereal* [10], a lightweight C++11 serialization library. We create a custom MPI derived datatype to encapsulate the serialized data in the send buffer for `MPI_Alltoallv`. Once the call completes, we deserialize the receive buffer to obtain the list of tuples, which contain the macro-nodes to be updated in 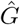. Lastly, we add the updated macro-nodes to a buffer (*rewire_list*); this is to initiate the rewiring of all modified macro-nodes.

### 9) Contig Generation: Gather and Walk

As described in Algorithm 3, at the end of the iterative phase, we are left with a total number of macro-nodes *<ψ* across all processes. At this stage, each process prepares a list of distinct starting points for initiating a walk in the compacted *PaK-Graph*. Entries in the *begin_kmer_list* are identified as the (*k*-1)-mer of macro-nodes with a terminal prefix (and *visit_count>* 0). Given that the graph has been sufficiently compacted (such that it can fit the memory of a single node), we initiate a call to `MPI_Allgatherv` to collate and gather all remaining macro nodes from all the processes. Thus each process now effectively receives a copy of compacted 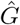, i.e., 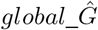.

The final phase of the contig generation algorithm involves the traversal (or walk) across the nodes in 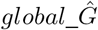. As described in Algorithm 4, we begin enumerating a contig for each entry in *begin_kmer_list*. Each MPI process initializes a contig and appends to it the terminal prefix extension followed by the macro-node, (*k*-1)-mer, and then initiates a walk, wherein it looks up its corresponding suffix extension in the wiring table and appends it to the contig. If the suffix extension is not terminal, the process continue the walk in a recursive fashion until a terminal suffix is encountered, at which time the walk is completed and the contig is returned as output. In the interest of space, we omit a detailed algorithmic pseudocode for the *walk* function.

#### Algorithm 4 Walk algorithm to generate final contigs

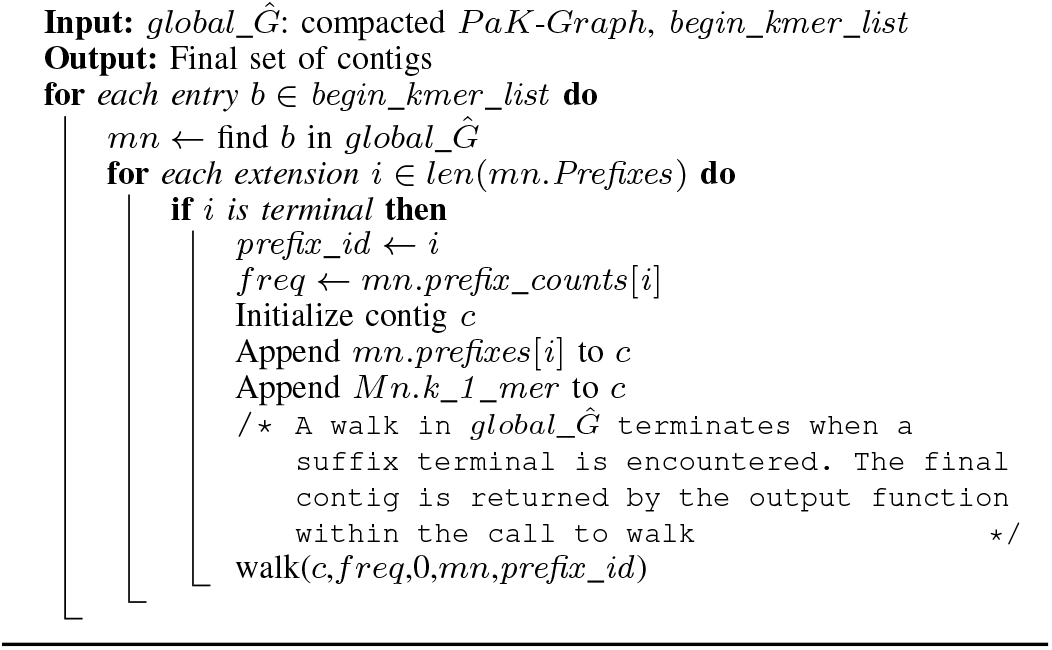

We illustrate the walking algorithm in Fig. 5, wherein we depict the contigs enumerated across three macro-nodes. The numbers on each wire represent the *visit_count* for the corresponding prefix/suffix extensions and the tuple (in brackets) indicates the: {offset in suffix, count} on the wire. The walking algorithm ensures that at no point during the walk, will the same sub-range in a wire be walked more than once. As seen in the case of contigs 1 and 2, the edge *GCATT* connecting the first and second node is traversed as part of both contigs. However, since they traverse separate ranges within the wire, both walks are disjoint and can occur concurrently.

**Fig. 5:**
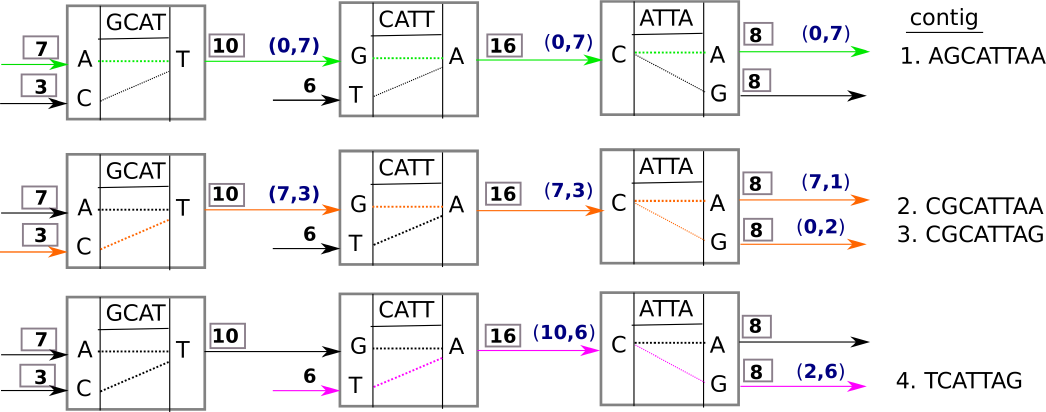
Walk algorithm illustration.

Lastly, we summarize the properties of the walking algorithm: a) Any walk will start at a terminal prefix and end at a terminal suffix; b) Every walk will terminate, and two walk’s starting from two different terminal prefixes will be guaranteed to be disjoint; c) There might be instances where a walk may traverse the same node multiple times owing to presence of repeat regions; and d) The algorithm does not guarantee that all repeat regions will be reachable from a terminal prefix, and thus covered as part of the contigs.

### D. Limitations of the PaKman algorithm

In this section, we discuss some of the implications of our algorithmic choices and point to avenues for further research:

- The performance of the *input reading* phase depends largely on the I/O subsystem. We have studied the impact of I/O related parameters across multiple filesystems and have presented our analysis in section IV-B.
- Our current implementation of *k-mer counting* does not account for overlapping communication and computation, and yet performs significantly better than the state-of-the-art. Once integrated, we hope to further improve the scalability of this phase.
- Our wiring algorithm relies on a greedy heuristic that takes the visit count of a *k*-mer into account while determining neighbors. As a result, we have observed instances wherein a false connection may lead to a mismatch, thereby, contributing to unaligned regions in the contigs. The quality of the wiring decisions made can be potentially improved with the incorporation of auxiliary information (such as paired-end) as available.

## IV. Experimental Evaluation

We evaluated *PaKman* using the datasets shown in Table I. All read datasets contain single-end reads generated from the real genome sequences, using the ART Illumina read simulator [11]. This approach is consistent with practice, as the simulators record the originating location for each read which can be later used as groundtruth for validation. The full human reference genome was obtained from the 2009 assembly (hg19, GRCh37 (GCA 000001405.1)). The bread wheat dataset (Triticum aestivum) corresponds to the GCA 900000045.1 Synthetic W7984 assembly and was obtained from NCBI (www.ncbi.nlm.nih.gov). For the purpose of extensive performance and qualitative evaluation, we used *C. elegans*, human chr 2+3, and full human data sets. The results of evaluating *PaKman* on the bread wheat genome are included in the Appendix.

**TABLE I:** Input datasets used in our experiments. *bp* refers to base pair.

| Organism | Genome size (bp) | Coverage | No. of reads | Read length | File size (GB) |
| --- | --- | --- | --- | --- | --- |
| *C.elegans* | 100,286,401 | 100x | 100,286,100 | 100 | 11 |
| human chr 2+3 | 441,221,803 | 100x | 432,999,339 | 100 | 58 |
| full human | 3,095,677,412 | 100x | 2,861,320,858 | 100 | 383 |
| bread wheat | 9,134,017,866 | 90x | 6,778,801,997 | 100 | 949 |

We used *k*=32 and *l*=8 for generating all *PaKman* results.

Our distributed memory experiments were conducted on the NERSC Cori machine (Cray XC40), where each node has 128GB DDR4 memory and is equipped with dual 16-core 2.3 GHz Intel Haswell processors. The nodes are interconnected with the Cray Aries network using a Dragonfly topology. Cori supports different file systems including: a) Lustre file system (with 30 PB of disk and > 700 GB/second I/O bandwidth), b) Burst Buffer, and c) GPFS. While we primarily focus our comparative evaluation on the Lustre file system, we detail *PaKman*’s I/O behavior on all three file systems. Our shared memory experiments were conducted on Keeran, a single-node machine with 750GB memory and dual 20-core 2.20 GHz Intel Xeon(R) CPU E5-2698 processors.

### A. Comparative Evaluation

#### 1) Evaluation on Distributed Memory System

We compared the performance of *PaKman* with the latest version of *HipMer* (v0.9.6.3.1), a state-of-the-art distributed memory genome assembly tool [12], [13]. We used the installation by the authors on the NERSC Cori system. Parameters for *HipMer* were left as default, with *k*=31. For *HipMer* we combine the execution times obtained from their log files corresponding to the tags ‘ufx-31’ and ‘meraculous-31’ and report that as the total time. Results were obtained with ppn=16 for both assemblers. Table IV shows Lustre striping details.

Fig. 6 presents strong scaling results for both assemblers for all three datasets. The plots show that both tools exhibit almost linear speedup under strong scaling. However, *PaKman* is considerably faster than *HipMer* on all inputs and processor sizes tested—by at least a factor of 2.2× (full human; no. cores=8K) and at most a factor of 3.2× (*C. elegans*; no. cores=32). *Using PaKman*, *we are able to assemble a complete set of contigs for the full human genome in 78.4 seconds on 8K cores*.

**Fig. 6:**
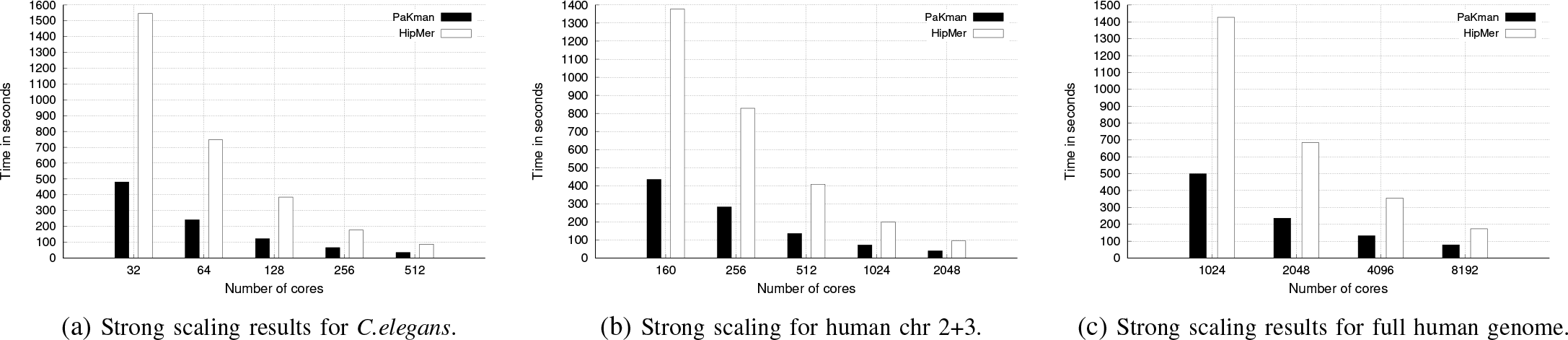
Strong scaling results for *PaKman* vs. *HipMer*

#### 2) Evaluation on Shared Memory System

Even though *PaKman* is designed for distributed memory machines, it can also be used on shared memory systems that support MPI. Consequently, we also performed a head-to-head comparison of *PaKman* running *p* MPI processes versus the shared memory tools running *p* threads on the same machine (Keeran). We compared against two state-of-the-art shared memory tools, namely *IDBA-UD* [1] and *FastEtch* [14].

Fig. 7 presents the results of the comparison for the *C.elegans* dataset. As can be seen, *PaKman* is the only tool to show near-linear scaling, while the remaining tools hardly show any improvement with number of cores. More importantly, for all core counts tested, we observe that *PaKman* is considerably faster than the other two tools; for instance, *PaKman* is faster than the second fastest tool which is IDBA-UD, by at least 2.8× (8 cores) and at most 9.3× (40 cores). These improvements are significant as these were observed despite the overheads associated with MPI.

**Fig. 7:**
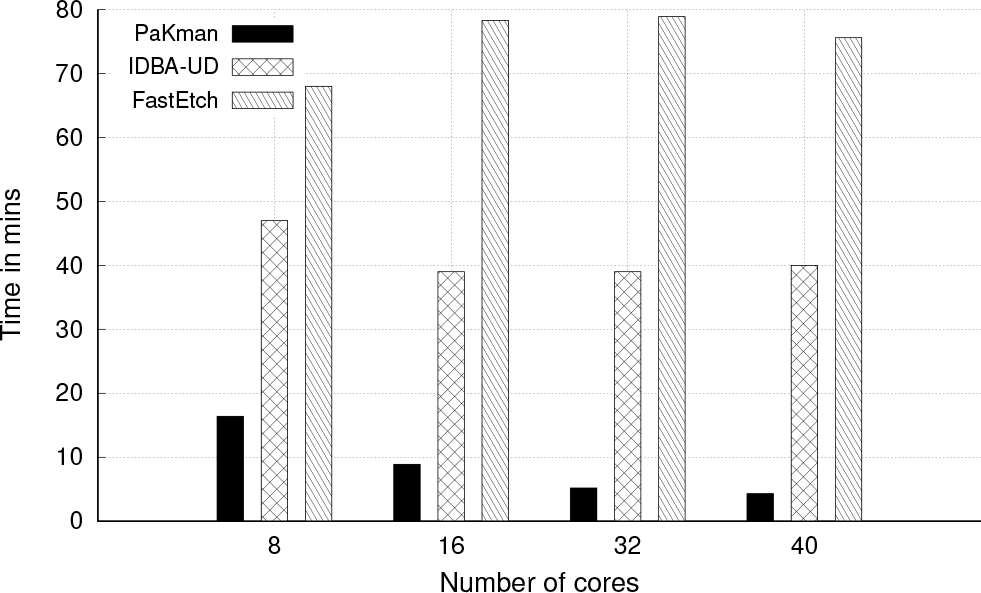
Single node shared memory scaling results for *C.elegans* across multiple assemblers

#### 3) Quality Evaluation

We compared the output quality of the assemblies produced by the various tools we tested. For this purpose, we compare the output contigs generated against the groundtruth (known genome from which reads were generated). The QUAST [15] tool was used for this comparison. The quality metrics reported are as follows: total number of contigs and the largest contig length; N50 contig length (larger the better); % of genome covered (larger the better); and largest alignment length (larger the better).

Table II summarizes our qualitative evaluation. As can be observed, *PaKman* generally outperforms or performs comparably to the second best tool by almost all metrics. We note here that, to enable a comparison, we did not test inputs with that paired-end read information, where an estimate of genomic distance is provided between pairs of reads (at input) alongside sequence information. This is because our tool does not yet include this feature; whereas tools such as *HipMer* and IDBA-UD do. We expect that with paired-end information these quality comparison results could change. Yet, without paired-end information, we observed *PaKman* to be competitive. It is also to be noted that *HipMer* proceeds more conservatively during the contig generation step and is known to produce longer contigs at the end of the scaffolding step.

**TABLE II:**
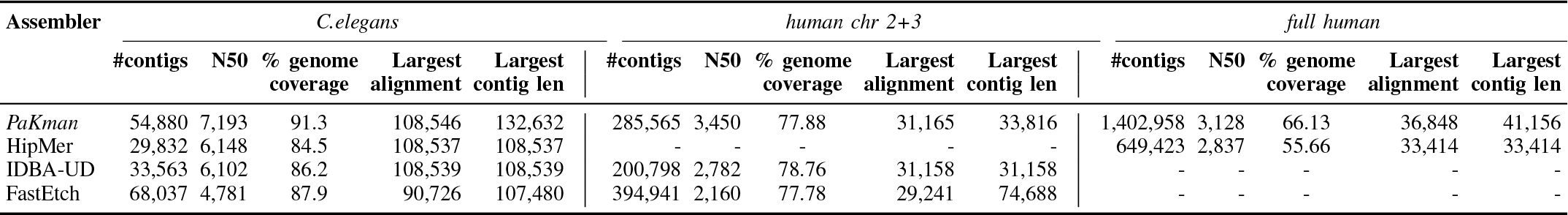
Quality statistics across assemblers. ‘-’ denotes a failed run (due to tool errors or exceeding memory capacity).

### B. PaKman: Detailed Performance Evaluation

To better understand the behavior of each phase we further break down the execution time. Fig. 8 presents the strong scaling results of *PaKman* broken down by its phases. The *input reading* step is I/O bound, dominated by calls to MPI-IO. For this step, while an improvement in time can be seen with increase in the number of cores, speedup is hardly linear. However, the runtime contribution to the total time is negligible (<1%). On the other hand, *kmer counting* is the most expensive step and that step scales linearly with the number of cores. The *contig generation* step, which includes most of the communication-intensive steps such as iterative graph compaction, also shows near linear speedup although for larger core sizes the speedup marginally deteriorates (as can be expected).

**Fig. 8:**
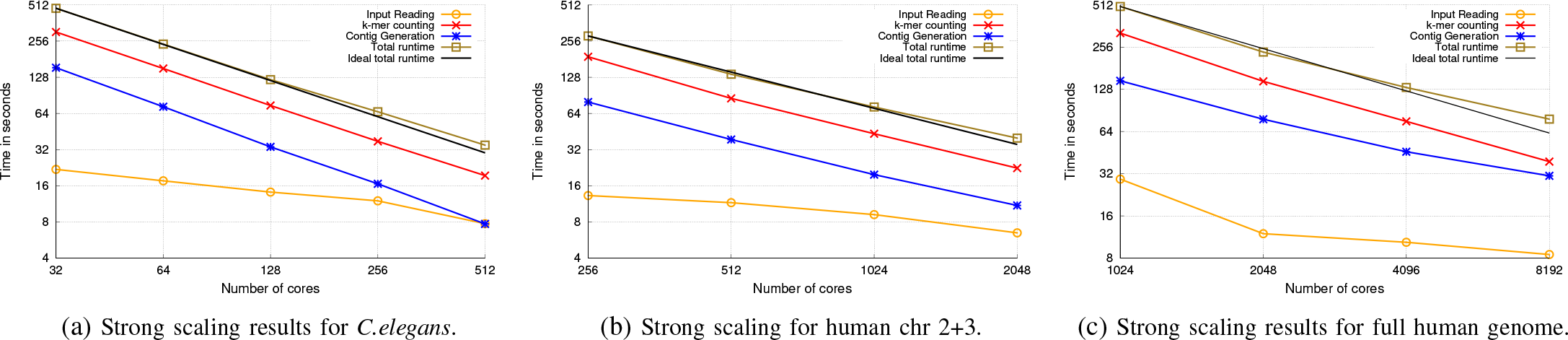
Strong scaling results for *PaKman* across multiple datasets. Total runtime corresponds to sum of all the phases.

Fig. 9 shows *PaKman*’s runtime broken down by computation, communication, and I/O. We observe that our algorithm scales efficiently and is noticeably compute bound, with the contribution from communication remaining under 20% even for large core counts. This makes the algorithm well-suited for massively parallel systems that offer greater support for compute operations than communication bandwidth. The current implementation used blocking collectives. With non-blocking collectives, there is the potential for further improvements.

**Fig. 9:**
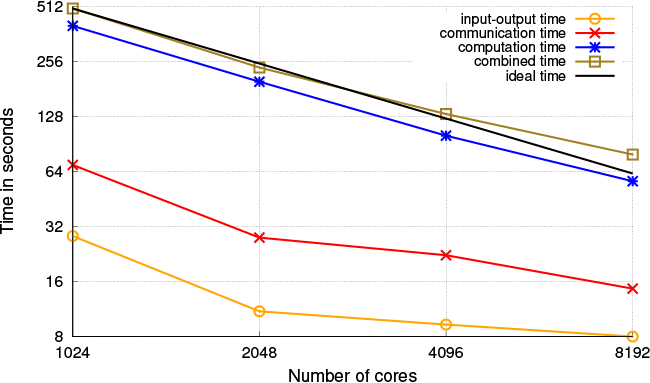
*PaKman* breakup of total time spent in I/O, communication and computation for full human genome

Fig. 10 captures the computation vs. communication break-down for the individual steps of *PaKman*. As can be expected, the contig generation step uses most of the communication time.

**Fig. 10:**
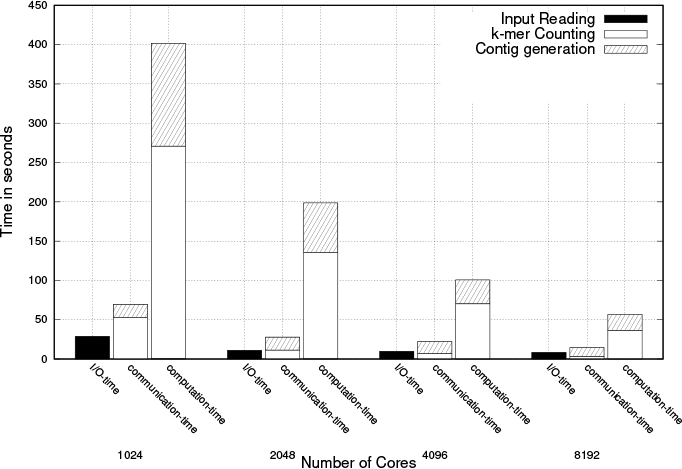
*PaKman* computation vs. communication time for various stages for full human genome

We note here that our algorithm uses only five distinct collectives (excluding the calls to MPI-IO), as shown in Table III. This is a useful property to have in MPI implementations, greatly simplifying performance portable installations on new parallel systems.

**TABLE III:** List of all MPI calls and their respective counts in each *PaK man* phase. Term ‘*b*’ in *k-mer Counting* phase denotes the number of batch rounds of communication; term ‘*i*’ in *Contig Generation* phase denotes the number of iterations of the *(de) serialize and transfer* phase in a given run

| PaKman Phase | MPI directive | #calls |
| --- | --- | --- |
| Input reading | MPI_File_open | 1 |
|  | MPI_File_get_size | 1 |
|  | MPI_File_read_at_all | 1 |
|  | MPI_File_close | 1 |
| k-mer counting | MPI_Allreduce | 2 |
|  | MPI_Alltoall | *b* *1 |
|  | MPI_Alltoallv | *b* *2 |
| Contig generation: MN-node construction | MPI_Alltoall | 1 |
|  | MPI_Alltoallv | 1 |
|  | MPI_Allreduce | 1 |
| Contig generation: (de)serialize and transfer | MPI_Allreduce | *i* *1 |
|  | MPI_Alltoall | *i* *2 |
|  | MPI_Alltoallv | *i* *1 |
| Contig generation: final gather | MPI_Allgather | 2 |
|  | MPI_Allgatherv | 1 |

As for the I/O time, Fig. 11 shows I/O scaling of the input reading step, for different file system configurations (Burst Buffer, Lustre, and GPFS). In the case of Burst Buffers (BB), we need to scale up the number of BB nodes with compute nodes, to keep the BB nodes busy but not over-subscribed. The performance was comparable to our Lustre runs. However, we needed to tune the BB settings for each run. While GPFS served well for the smaller datasets, for the full human genome, it did not scale beyond 2048 processes.

**Fig. 11:**
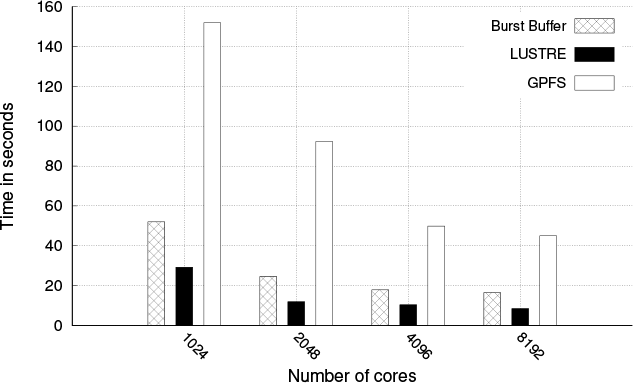
I/O scaling of the Input Reading phase of *PaKman* for the full human dataset across various file systems.

Table IV shows the Lustre settings used for our *PaKman* runs. We observed that, unlike *HipMer*, the total time for *PaKman* responded to changes made to the striping configuration; this configuration can be varied and tested quickly for improving *PaKman*’s performance. Furthermore, we observe that Lustre I/O times (for the settings presented) remain constant for a given dataset across all our experiments. Table IV also lists various other statistics for *PaKman*. We observed that the number of macro-nodes in the initial PaK-Graph is roughly an order of magnitude smaller than the number of distinct *k*-mers. This result shows the initial degree of compression that *PaKman* achieves (even before graph compaction) compared to standard de Bruijn graph implementations.

**TABLE IV:** *PaKman* runtime statistics. Lustre parameters: *sc* denotes stripe count, *ss* denotes stripe size; *cb* denotes the number of MPI aggregators (*cb_nodes*).

| **PaKman** | *C.elegans*<br>(**p=512**) | **chr 2+3**<br>(**p=2048**) | **full human**<br>(**p=8192**) |
| --- | --- | --- | --- |
| # distinct k-mers | 1,585,416,564 | 6,619,495,237 | 42,182,152,288 |
| #total Macro nodes | 188,188,574 | 793,197,434 | 4,987,035,369 |
| #total MPI calls | 155 | 1367 | 2847 |
| Total time (secs) | 34.87 | 39.88 | 78.42 |
| Lustre I/O parameters | sc=25, ss=4M,<br>cb=16 | sc=50, ss=8M,<br>cb=64 | sc=50, ss=8M,<br>cb=256 |

Because a significant fraction of contig generation phase is spent in communication, we take a deeper look at the performance breakdown of the individual steps within the phase. Shown in Fig. 12, the steps *Initial setup wiring* and *Generate Independent set* scale linearly and are almost negligible on 8K cores run. In spite of being communication-intensive, the *Iterate and pack nodes* step also scales linearly. Scalability is limited for the *(de)serialize and transfer*, as it is highly communication-bound during iterative graph compaction. The amount of communication involved in *Macro node construction* and *Gather and Walk* is minimal and does not impact the overall performance.

**Fig. 12:**
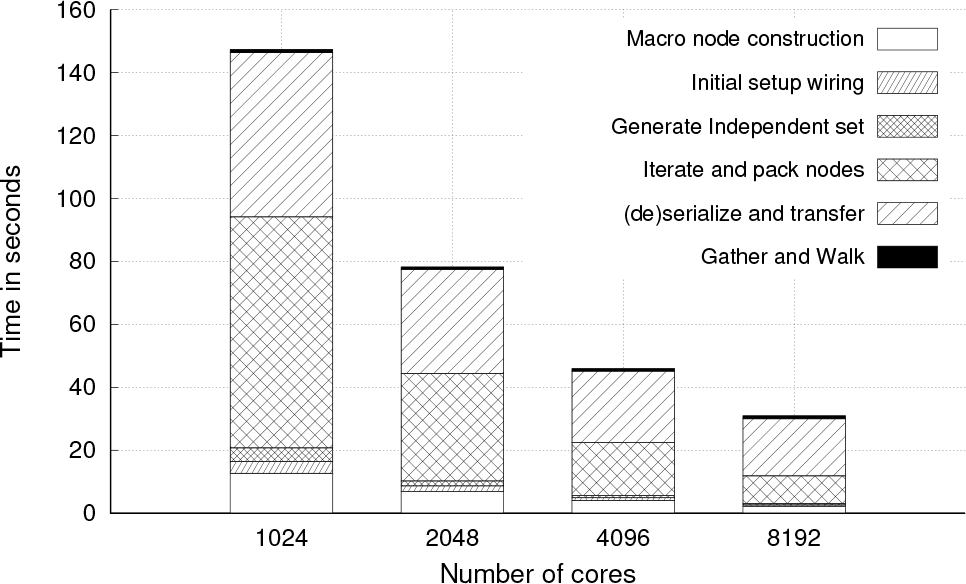
*PaKman* performance breakup of the several stages in the *contig generation* step for the full human genome.

Fig. 13 shows the performance breakdown of first 40 (out of 708) iterations on the iterative phase of the contig generation step on 1K cores for the full human genome. We observe a superlinear decrease in total time, which plateaus after the first 20 iterations for both plots. Fig. 13a also shows the number of macro-nodes and the independent set size, at each iteration. We observe that the fraction of macro-nodes included in the independent set size at each iteration shrinks gradually from 28% to 13% at the end of 40 iterations, to eventually 7% in the final iteration. Fig. 13b shows that the contributions to runtime vastly diminish after the first 10-12 iterations.

**Fig. 13:**
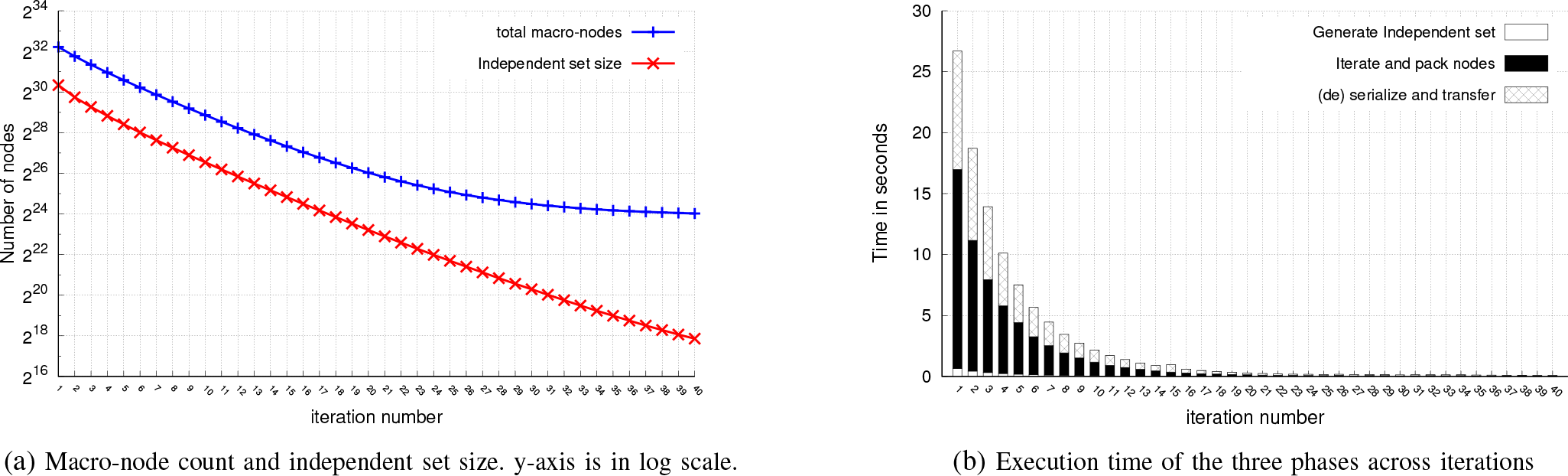
*PaKman* behavior of the first 40 iterations in iterative phase of contig generation for full human genome (p=1024).

### C. Parametric evaluation

We evaluated the effect of varying two input-based parameters—viz., coverage and read length—on the performance of *PaKman*. Table V presents the statistics across all four full human datasets, wherein we compare the baseline dataset with three datasets of the same genome with varying read length and coverage.

**TABLE V:**
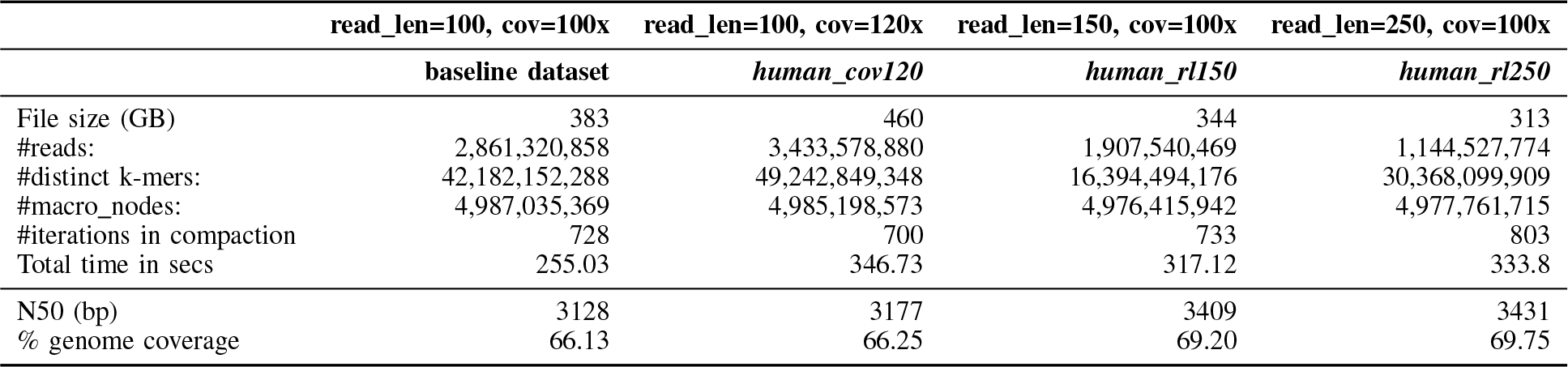
*PaKman* performance evaluation across four full human datasets with varying parameters on 2048 processes.

As seen in Table V, increasing the coverage or the read length causes an increase in total runtime. Specifically, an increase in coverage contributes to a larger number of reads, which subsequently increases the work during the *k-mer counting* phase. We notice a similar effect for increasing the read length, wherein despite the presence of fewer reads, more work is needed to parse each read given its longer length, to generate the *k*-mers.

While these results are to be expected, we also observed that despite the increase in number of distinct *k*-mers, the number of macro-nodes resulting from our threshold-based pruning step (post-*k-mer counting*) remains relatively uniform across the different input settings. This is a desirable property owing to two reasons: a) Not increasing the number of macro-nodes implies negligible impact on the *contig generation* time as seen in Fig. 14a; b) Secondly, note that the macro-nodes capture the key *information* encoded within the input reads that contribute toward the output contigs; therefore these results show the ability of our pruning step to capture that information in a stable manner even as the input read length or coverage changes. Eventually, after contig generation, we see the positive impact of increasing coverage and read length on the output quality metrics (N50, coverage). We also note that the quality improvement is better with increased read length than with increased coverage. This is to be expected as longer reads (with a lower error rate) are a more valuable source of information than increased coverage (which simply increases the redundancy in information beyond a certain value).

**Fig. 14:**
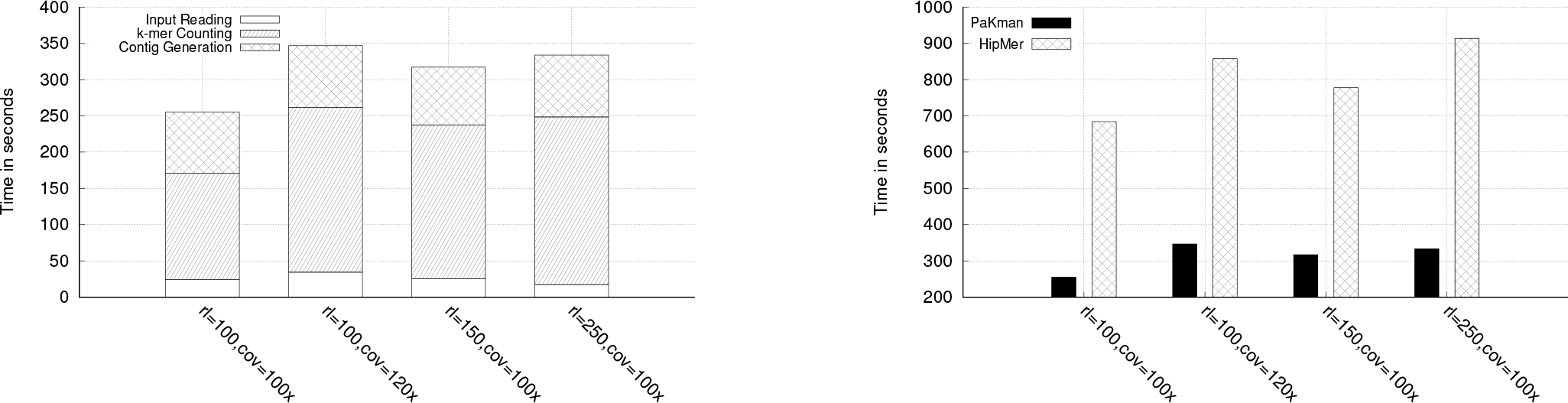
*PaKman* Performance across all four full human datasets with varying parameters for p=2048. *rl* corresponds to read length and *cov* corresponds to coverage.

The effects of these two parameters on *HipMer*’s performance were similar to that of *PaKman* as shown in the Fig. 14b. In terms of quality, *HipMer* presented a marginal increase in N50 (from 2837 to 2843) with the increase in read length.

## V. Related Work

De novo genome assembly is a widely researched topic with a number of assembly tools and algorithms developed over the last two decades. Therefore in this section, we focus primarily on parallel short-read assemblers. Short read assemblers correspond to that subset which target reads generated from NGS technologies (e.g., Illumina, 454 pyrosequencing, SOLiD). Most modern day short read assemblers have widely adopted the de Bruijn Graph-based (DBG) method of assembly, originally introduced by Pevzner et al. [16]. Popular shared memory DBG based assemblers include (but not limited to): Velvet [2], ALLPATHS-LG [17], and SOAP-denovo [3], all of which utilize OpenMP/Pthreads for parallelization. More recent implementations of the method include IDBA-UD [1], an iterative DBG assembler that generates assemblies by sequentially iterating from small to large *k*-values used in graph construction. Although OpenMP parallelized (for a single *k*), this method can be time intensive since graphs for multiple *k* values proceed sequentially. SPAdes [5] has support for multithreading and produces assemblies of high quality owing to its detailed error correction step. However, it is costly with respect to the amount of time and memory it consumes. We were unable to run SPAdes for our medium to large datasets owing to its significant memory footprint.

Notable examples of distributed memory DBG based assemblers include Ray [18], PASHA [19] and YAGA [20]. Ray and YAGA have shown to be scalable except for the I/O that proves to be a bottleneck when reading and writing to files. ABySS [4] is a full end-to-end assembler and is one the first to be parallelized using MPI and was the first software to assemble a human genome from short reads. However their input reading step presents a bottleneck to the overall performance. ABySS 2.0 [21] departs from using MPI and instead employs Bloom filters to represent a de Bruijn graph and reduce memory requirements. HipMer [13], [12] as discussed in the previous section, also uses Bloom filters to generate its version of the de Bruijn graph. In our evaluation, we observed HipMer to scale well for all sizes of input data at high core counts. SWAP [22] and SWAP-2 [23] are also among the newer set of assemblers that have been MPI parallelized for executing at large scale for large genomes. Although the assembly output from SWAP-2 was high in quality, we observed it failed to execute on NERSC Cori beyond small to medium sized datasets.

## VI. Conclusions

We introduced *PaKman*, a new algorithm for efficient scaling of two crucial phases of the genome assembly pipeline. We presented a new data structure—*PaK-Graph*—for contig generation with simplified communication requirements. Our method demonstrated a speedup of over 2× on average in comparison with state-of-the-art distributed and shared memory implementations, respectively.

Our goals for future work include: a) Incorporation of paired end information toward improving quality: although not trivial the wiring function has the capacity to be extended for this purpose. But a space-efficient representation will be needed to make it work as the graph compacts. b) Provide an end-to-end solution, by incorporating the scaffolding phase to fully complete the assembly pipeline. c) Extension to scenarios where reads are of variable lengths (possibly even long reads). d) Extension to use heterogeneous architectures.

**Fig. 15:**
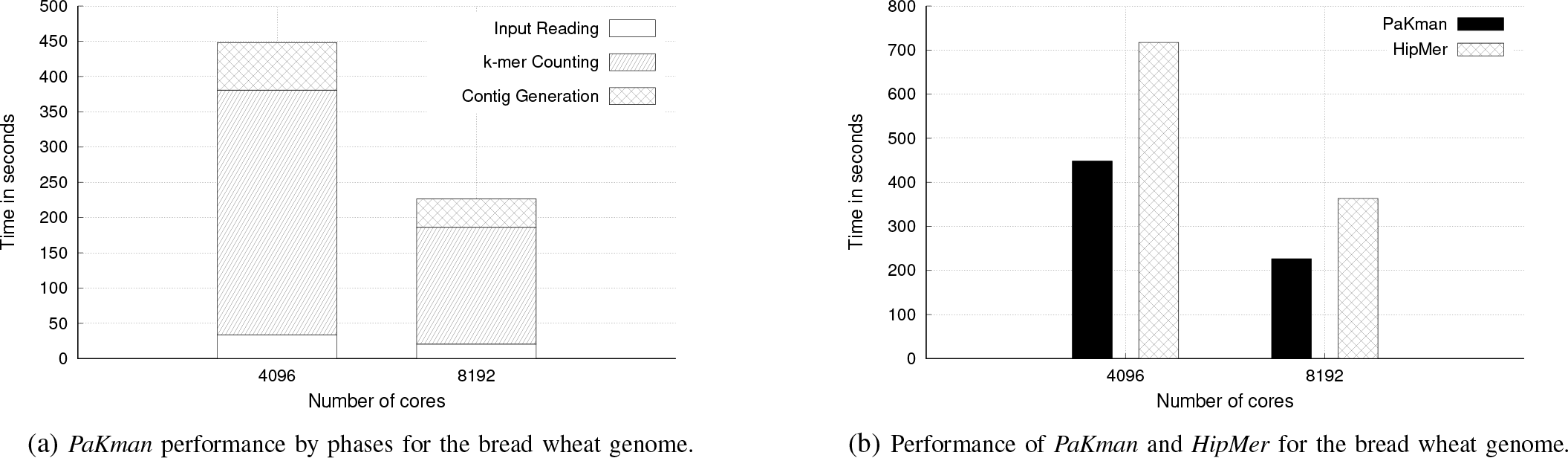
Performance evaluation for the bread wheat genome.

## APPENDIX A PaKman PERFORMANCE EVALUATION FOR PLANT GENOME

We also conducted experiments on a larger more complex genome namely the bread wheat genome, characterized as highly repetitive and much larger in size (more than three times the size of the full human genome). The bread wheat genome utilized in our experiments spans 9.1Gbp (over 56%) of the 16Gbp genome of hexaploid wheat, *Triticum aestivum*.

Fig. 15 presents strong scaling scaling results for both *PaKman* and *HipMer* for the bread wheat dataset. Fig. 15a presents the breakdown in time for all the distinct phases of *PaKman*. Fig. 15b shows the total runtime wherein we observe that both tools exhibit almost linear speedup. *PaKman* completes execution of the bread wheat dataset in 226 seconds on 8K cores and is reported as 1.6x faster than *HipMer*.

## References

[1] Y. Peng, H. C. Leung, S.-M. Yiu, and F. Y. Chin, “Idba-ud: a de novo assembler for single-cell and metagenomic sequencing data with highly uneven depth,” Bioinformatics, vol. 28, no. 11, pp. 1420–1428, 2012.

[2] D. Zerbino and E. Birney, “Velvet: algorithms for de novo short read assembly using de bruijn graphs,” Genome research, pp. gr–074 492, 2008.

[3] R. Li, H. Zhu, J. Ruan, W. Qian, X. Fang, Z. Shi, Y. Li, S. Li, G. Shan, K. Kristiansen et al., “De novo assembly of human genomes with massively parallel short read sequencing,” Genome research, vol. 20, no. 2, pp. 265–272, 2010.

[4] J. T. Simpson, K. Wong, S. D. Jackman, J. E. Schein, S. J. Jones, and I. Birol, “Abyss: a parallel assembler for short read sequence data,” Genome research, pp. gr–089 532, 2009.

[5] A. Bankevich, S. Nurk, D. Antipov, A. A. Gurevich, M. Dvorkin et al., “SPAdes: a new genome assembly algorithm and its applications to single-cell sequencing,” J. Comp. Bio., vol. 19, no. 5, pp. 455–477, 2012.

[6] E. W. Myers, “The fragment assembly string graph,” Bioinformatics, vol. 21, no. suppl 2, pp. ii79–ii85, 2005.

[7] A. Kalyanaraman, S. J. Emrich, P. S. Schnable, and S. Aluru, “Assembling genomes on large-scale parallel computers,” in IPDPS, 2006, pp. 10–pp.

[8] R. Chikhi, A. Limasset, S. Jackman, J. T. Simpson, and P. Medvedev, “On the representation of de bruijn graphs,” in International conference on Research in computational molecular biology, 2014, pp. 35–55.

[9] E. Cohen, “Min-hash sketches,” Encyclopedia of Algorithms, pp. 1–7, 2008.

[10] W. S. Grant and R. Voorhies, “cereal–a c++ 11 library for serialization,” URL https://github.com/USCiLab/cereal, 2013.

[11] W. Huang, L. Li, J. R. Myers, and G. T. Marth, “Art: a next-generation sequencing read simulator,” Bioinformatics, vol. 28, no. 4, pp. 593–594, 2011.

[12] E. Georganas, A. Buluç, J. Chapman, L. Oliker, D. Rokhsar, and K. Yelick, “Parallel de bruijn graph construction and traversal for de novo genome assembly,” in SC, 2014, pp. 437–448.

[13] E. Georganas, A. Buluç, J. Chapman, S. Hofmeyr, C. Aluru, R. Egan, L. Oliker, D. Rokhsar, and K. Yelick, “Hipmer: an extreme-scale de novo genome assembler,” in SC, 2015, p. 14.

[14] P. Ghosh and A. Kalyanaraman, “Fastetch: A fast sketch-based assembler for genomes,” IEEE/ACM TCBB, 2017.

[15] A. Gurevich, V. Saveliev, N. Vyahhi, and G. Tesler, “Quast: quality assessment tool for genome assemblies,” Bioinformatics, vol. 29, no. 8, pp. 1072–1075, 2013.

[16] P. A. Pevzner, H. Tang, and M. S. Waterman, “An eulerian path approach to dna fragment assembly,” PNAS, vol. 98, no. 17, pp. 9748–9753, 2001.

[17] S. Gnerre, I. MacCallum, D. Przybylski, F. J. Ribeiro, J. N. Burton, B. J. Walker, T. Sharpe, G. Hall, T. P. Shea, S. Sykes et al., “High-quality draft assemblies of mammalian genomes from massively parallel sequence data,” PNAS, vol. 108, no. 4, pp. 1513–1518, 2011.

[18] S. Boisvert, F. Laviolette, and J. Corbeil, “Ray: simultaneous assembly of reads from a mix of high-throughput sequencing technologies,” Journal of computational biology, vol. 17, no. 11, pp. 1519–1533, 2010.

[19] Y. Liu, B. Schmidt, and D. L. Maskell, “Parallelized short read assembly of large genomes using de bruijn graphs,” BMC bioinformatics, vol. 12, no. 1, p. 354, 2011.

[20] B. G. Jackson, M. Regennitter, X. Yang, P. S. Schnable, and S. Aluru, “Parallel de novo assembly of large genomes from high-throughput short reads,” in IPDPS, 2010, pp. 1–10.

[21] S. D. Jackman, B. P. Vandervalk, H. Mohamadi, J. Chu, S. Yeo et al., “Abyss 2.0: resource-efficient assembly of large genomes using a bloom filter,” Genome research, pp. gr–214 346, 2017.

[22] J. Meng, B. Wang, Y. Wei, S. Feng, and P. Balaji, “Swap-assembler: scalable and efficient genome assembly towards thousands of cores,” in BMC bioinformatics, vol. 15, no. 9. BioMed Central, 2014, p. S2.

[23] J. Meng, S. Seo, P. Balaji, Y. Wei, B. Wang, and S. Feng, “Swap-assembler 2: Optimization of de novo genome assembler at extreme scale,” in ICPP. IEEE, 2016, pp. 195–204.

